# Dual immunosensor based on methylene blue-electroadsorbed graphene oxide for rapid detection of influenza virus antigen

**DOI:** 10.1101/045310

**Authors:** Murugan Veerapandian, Robert Hunter, Suresh Neethirajan

## Abstract

Rapid detection of influenza viral infections in the poultry farm is advantageous in several facts such as environmental/personal safety, food-security and socio-economy. Herein, we report the development of an electrochemical based dual sensor platform composed of methylene blue-electroadsorbed graphene oxide nanostructures modified with monoclonal antibodies, H5N1 and H1N1. Bio-functional layers comprised of chitosan and protein-A molecules were implemented at the interface of sensor element and antibodies, which synergistically enriched the bio-activity of immobilized antibodies for the immune complex formation. The differential pulse voltammetric signals resulted from the developed immunosensor platform exhibited a good correlation (R^2^= 0.9978 for H1N1 and R^2^=0.9997 for H5N1) for the wide range of target concentrations (25 to 500 pM). Chronoamperometric study also revealed the amplified current sensitivity of the immunoelectrodes even at the picomolar level. The proposed immunosensor design not only provide rapid analytical response time (<1 min) but also provide simplicity in fabrication and instrumentation which paves an attractive platform for on-farm monitoring of viral infections.

## 1. Introduction

Infectious influenza A virus pathogen that affect poultry can be categorized into two groups based on the virulence of the disease, high-pathogenic and low-pathogenic avian influenza (AI) viruses [1]. Among the high-pathogenic AI strains, the subtype H5N1 is often responsible for the systemic infection which results in socio-economic losses around the world [2]. A recent report identified an estimated loss of $309.9 million, in poultry production and related businesses, in greater Minnesota [3]. After avian flu outbreak reported in December 2014, the government of Canada is progressively strengthening the budget on the surveillance, early detection and response measures to avian flu. Transmission of high pathogenic virus strains from poultry facility to healthy environment could lead serious pandemics. Public health agency of Canada documented a seasonal flu (in humans) cases of approximately 12,200 hospitalizations and, on average, 3,500 deaths each year [4]. A global alert to AI virus infection due to the subtypes of H5N1 to H7N9 has been well documented in the literature [5]. In addition to vaccination, early and accurate detection of virus pathogen is crucial for the prevention of disease transmission. At the moment, there is no on-farm diagnosis available for the rapid detection of AI pathogen. Most of the time the analyte samples collected from throat of live animals, fecal content and blood components from the poultry location is transported to the centralized laboratory for analysis.

Virus isolation culture is the conventional standard widely practiced for the detection of influenza strains. But this technique requires intensive labor and time consuming procedures. Enzyme-Linked Immunosorbent Assay (ELISA) and other molecular methods like PCR and RT-PCR are also used due to the better sensitivity and less operational time, compared with traditional methods [6]. However, limitations associated with these techniques are necessity of trained personnel to isolate genetic material and need of expensive instrumental facilities, which are not suitable for on-farm monitoring and small diagnostic laboratories. In the recent past, biosensor for influenza virus based on DNA-aptamer immobilized on Surface Plasmon Resonance (SPR) active element [7], microfluidic chip supported with magnetic bead and labeled quantum dots [8], and colorimetric assay based on enzyme-induced metallization [9] are demonstrated. However, these techniques are still depended on experienced technician and have issues in the field deployability. Recent advancements in bio-diagnostics desire a new technique with features of cost-efficient fabrication and portability, high sensitivity and integration feasibility for modern gadgets with user-friendly operation. Among various recent diagnostics, electrochemical biosensor has been investigated to be promising for sensitive detection of AI viral strains [6]. Due to its ease in miniaturization, simple instrumentation, rapid analysis time and user-friendliness, electrochemical sensors are advantageous for point-of-care application. Particularly, sensor surface based on electroactive nanomaterials supported affinity reaction of antigen-antibody are demonstrated to have greater sensitivity [10–12].

Electrochemical impedance spectroscopy based immunosensing platform have been studied on the surface of modified gold electrodes for the detection of influenza virus pathogen [13–15]. However, still optimization related to disposable redox-active electrode materials with durable bio-affinity functionalization and feasibility for dual or multiplexed sensor platform is always desired. Further, compared to conventional (disintegrated) three-electrode system, cost-efficient screen-printing technology derived electrodes are potential for compact device. Varieties of patterned electrodes are emerged as biosensor platform because of simple surface modification and integration [16–18]. In addition to bio-affinity, significant efforts have been devoted to amplify the electron-transfer properties of sensor element at the analyte-substrate interface [17]. In this context, biosensor platform based on two-dimensional nanostructure, graphene oxide (GO) modified hierarchical materials are received recent attention due to its cost-efficient precursor source and flexibility in the surface anchoring of redox species and bio-recognition molecules (viz., antibody, enzyme, DNA or RNA oligonucleotide, peptide) [19–21]. Owing to its abundant polar oxygen functional groups graphene oxide (GO) yield good dispersiblity in various solvents including water. Therefore, GO-dispersion can be subsequently coated on various solid substrates to prepare thin conductive films using drop-casting, ink-jet printing or spin coating and processed for electrode materials [19]. Further, the physicochemical properties of GO sheets can be surface engineered by chemical or biomolecular doping (*via* covalent or non-covalent reaction), photo-irradiation and thermal treatment [11, 19–21].

Considering the merits of screen printed electrodes (SPEs) and novel functions of redox active GO-derivatives toward biosensor application, herein a dual immunosensor platform based on methylene blue (MB) electroadsorbed GO has been designed for the first time to detect influenza virus antigen. It is well known that MB is a positively charged organic dye molecule with enhanced electrocatalytic properties. GO is known for multiple negatively charged oxygenated functional groups on its basal plane and edges (viz., carboxylic acid, epoxy, carbonyl, and hydroxyl), which are useful in electrostatic interactions. Surface integration of bio-active interface layers, such as chitosan (CS) and protein-A molecules, on sensor element are expected to have retained bio-activity of immobilized monoclonal antibodies. For proof-of-concept, two pathogenic influenza viral strains (H5N1 and H1N1) have been selected to test the immunosensing ability of the proposed system. Mechanism behind the formation of electroadsorbed sensor element, immobilization of bio-affinity layers and immunocomplex reaction at the electrode interface are comprehensively presented. Based on the fundamental voltammetric technique optimal applied potential suitable for chronoamperometric experiments are investigated. Present experimental study on MB-electroadsorbed GO based immunosensor platform enable sensitive electron transfer process for dual sensor application, which has the potential for on-farm simultaneous detection of pathogenic viral strains.

## 2. Experimental Section

### 2.1 Materials

Graphite powder (<20 μm, synthetic), methylene blue (dye content ≥82%), chitosan (low molecular weight: 50,000-190,000 g mol^-1^; 75-85% deacetylated), protein-A from *Staphylococcus aureus* and phosphate buffered saline (PBS) were purchased from Sigma-Aldrich. Mouse anti influenza-A hemagglutinin monoclonal H5N1 antibody, mouse anti influenza-A hemagglutinin monoclonal H1N1 antibody, recombinant hemagglutinin influenza A virus H5N1 Vietnam 1203/04 and recombinant hemagglutinin-influenza A virus H1N1 California 04/2009 were purchased from Cedarlane Labs, Ontario, Canada. Other chemicals were analytical standard and used as received without further purification. All experimental solutions were prepared using deionized (DI) water from Millipore system with resistivity >18.2 MΩ.

### 2.2 Electrodes

Disposable dual and single carbon screen-printed electrodes (SPEs) were customized from Dropsens, Llanera (Asturias), Spain, distributed by Metrohm Canada. The single electrochemical cell substrate (DRP-C110) consists of central circular carbon as working electrode (area 4 mm in diameter), crescent-shaped carbon as counter and silver as reference electrode. The dual electrode (DRP-C1110) have two elliptical carbon working (6.3 mm^2^ each) surface, with a common carbon counter electrode and a silver reference electrode, all screen printed on a ceramic substrate, (3.4 × 1.0 × 0.05 cm, length × width × height). Other single carbon-SPEs were customized from Pine research instrumentation, NC, USA. This consists of a carbon substrate with an area of 2 mm in diameter as working electrode, integrated U-shaped carbon and small circular Ag/AgCl as counter and reference electrodes, respectively. Overall card dimensions are 6.1 × 1.5 × 0.036 cm.

### 2.3 Preparation of modified electrodes

After performing initial rinse with DI water, the desired single or dual electrode’s carbon working surface was modified by drop casting 2 μL of aqueous GO dispersion (1 mg mL^-1^) and allowed to evaporate at (laboratory) room temperature for 15-20 min. Colloidal dispersions of GO nanosheet are prepared by harsh oxidation of graphite powder using modified Hummers’ method [22]. To ensure uniform modification on the working surface typically three layers (total) of casting were performed. As-fabricated GO-modified electrodes were then utilized for electrochemical adsorption of MB. The amount of MB adsorption was controlled by cyclic voltammetric technique (20 potential cycles) in the region □1.0 to +1.0 V, at a scan rate of 100 mV/s. For this, 10 mM PBS as electrolyte solution with an optimal concentration of 50 μg mL^-1^ MB was prepared. All modified electrodes were thoroughly rinsed with DI water for further experiments.

### 2.4 Immunosensor construction

Immunosensor fabrication was achieved in the support of bio-active interface layers. At first, two layers of chitosan (CS) (each 3 μL, 1 mg mL^-1^, prepared in 1% (w/v) acetic acid) was drop casted on to the GO-MB electrode surface and incubated at room temperature for 15-20 min. Afterward, 2 μL of protein A from *S. aureus* (50 μg mL^-1^ in PBS pH 7.4) was drop casted onto the GO-MB/CS electrode surface and incubated at room temperature for 45 min. At the end of each interface layer coating, electrodes were gently immersed in DI water and PBS solution to remove unbound CS and protein A molecules, respectively. Immobilization of the monoclonal antibodies (H5N1 or H1N1) was achieved by drop casting the antibodies (10 μg mL^-1^, 4 μL) onto the protein A-modified GO-MB/CS surface and allowing them to incubate for 45 min. After washing with buffer to remove surplus antibodies, the antibodies-modified electrode was treated with BSA solution (2 μL, 0.25% w/v), as blocking agent and incubated for 30 min to avoid non-specific interactions. After washing gently in buffer solution (5 sec), as-prepared immunosensor was stored at 4□8°C for further studies.

### 2.5. Immunodetection of H5N1 or H1N1 protein

H5N1 or H1N1 virus protein detection was based on antigen-antibody immune reaction. The recombinant H5N1 or H1N1 virus antigens were captured by the monoclonal antibodies immobilized on the protein A-modified GO-MB/CS electrode, which resulted in a change of peak current intensity. Desired analyte concentrations of recombinant H5N1 or H1N1 proteins prepared in PBS solution were dropped onto the electrode surface. Upon 30 min of incubation time, electrochemical measurements were performed to monitor the change of current (differential pulse voltammetric or chronoamperometric) response from the prepared electrodes. The typical sample volume requires to cover the integrated Dropsens electrodes were ~50 and ~100 μL for single and dual sensor surface, respectively. For electrodes customized from Pine research instrumentation requires ~300 μL.

### 2.6 Instrumentation

Surface topography of different material’s modified electrodes were studied from FEI Inspect S50, at an acceleration voltage of 15 V. Elemental distribution of the sensor surface was investigated using Oxford X-Max20 silicon drift detector and the data were processed by Aztec software. Electrochemical measurements were carried out with μStat 400 Biopotentiostat from DropSens, Llanera (Asturias), Spain. Data were analyzed using Dropview 8400 software. Cyclic voltammograms (CVs) were recorded in the potential window of □0.5 to □0.1 V, at a scan rate of 50 mV s^-1^ in 10 mM PBS (pH 7.4). The differential pulse voltammetric (DPV) measurements were screened in the potential region from □0.5 to □0.1 V at a scan rate of 50 mV s^-1^ with amplitude of 2 mV and a step potential of 4 mV. Chronoamperometric responses were studied at an applied potential of □0.275 V. Current densities specifically at 50 seconds were calibrated for the quantification of virus antigen concentration. As-measured electrochemical data were processed in OriginPro 8.5 (OriginLab Corporation, MA, USA) for the preparation of graphs.

## 3. Results and discussion

### 3.1 Immunoelectrode fabrication and working principle

Pictorial illustration shown in Scheme 1 describes the typical dual SPE and the sequence of surface modification employed in the development of immunosensor platform for two different AI antigens. Individual components exist on the sensor platform has their unique functionalities. For instance, thin atomic layers of GO with abundant oxygenated functional groups on basal planes and edges provide necessary chemical interaction sites to the redox molecules, in this case MB. As described in literature, chemically altered surface topography of GO exhibit modified configuration of sp^2^ and sp^3^ carbon domains, which offer new lattice orientation with numerous electroactive charge carriers suitable for sensor application [11]. MB as a redox probe is also known for multiple sensor studies. Owing to its anionic nature, GO are studied to have efficient adsorbent role to various cationic dyes, in particular concentration-dependent aggregation of MB on the GO surface lead to stable MB dimers and trimers for fluorescent quenching properties [23]. Unlike physical adsorption, electrochemically adsorbed MB molecules on graphene sheets are demonstrated to have enhanced electron-transfer properties for biomolecular detection [24]. Similarly, in this work electroadsorption of MB molecules on the surface of GO-modified electrodes are controlled by cyclic voltammetry. Biopolymer derivative, CS is widely employed as an interface layer for sensor platform. Due its intrinsic oxygen- and nitrogen-based functional groups CS is well suited for various covalent and/or cross-linking reactions. Further, excellent solubility in mild acidic aqueous solution, film-forming ability and electroconductivity of CS offers multifunctional role in biosensor design, reviewed elsewhere [25]. Protein-A with molecular weight 40□60 KDa is a well-known Fc receptor for the specific binding of antibodies (Fc region) [26]. Decoration of protein-A moieties on the sensor element promote the immobilization of antibodies, which results in the exposure of Fab active sites to epitopes, the selectivity of electrode toward the target antigens are strongly improved [27]. Working principle behind the proposed sensor platform is based on the complex formation between the immobilized monoclonal antibodies (mAb) and selective antigen samples. Such immunocomplex formation on the sensor element influences the inherent voltammetric response. Changes in the current response of surface modified electrodes before and after interaction with various concentrations of analyte samples were appropriately calibrated to determine the sensing parameters.

**Scheme 1.**
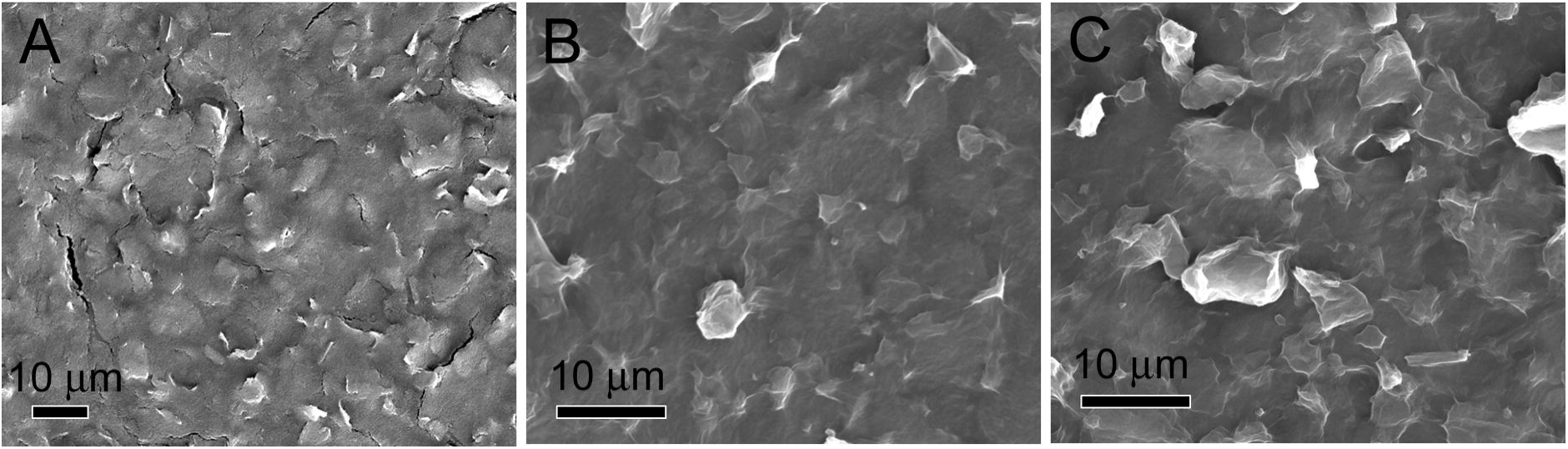
Representation of dual SPE and sequence of surface modification in the preparation of immunosensor platform. WE: working electrode, CE: counter electrode, and RE: reference electrode.

### 3.2 Surface morphology and Raman spectrocopy of electrode materials

Morphological features of pristine and GO-based materials modified on carbon SPEs (CSPEs) were studied from SEM (Fig. 1). Compared to pristine CSPE, surface of GO nanosheets-modified electrode is free from micro cracks. Existence of larger continuous network of stacked GO thin sheets on the working substrate allows better electroadsorption of MB molecules. Elemental composition on the surface of individual electrodes is studied by energy dispersive X-ray spectroscopy analysis in SEM (Fig. S1–S3). It can be observed that unlike pristine CSPE and GO-modified CSPE, surface of MB-electroadsorbed GO electrodes show additional elemental composition (S and N) associated to inherent MB, supporting the successful functionalization.

**Fig. 1.**
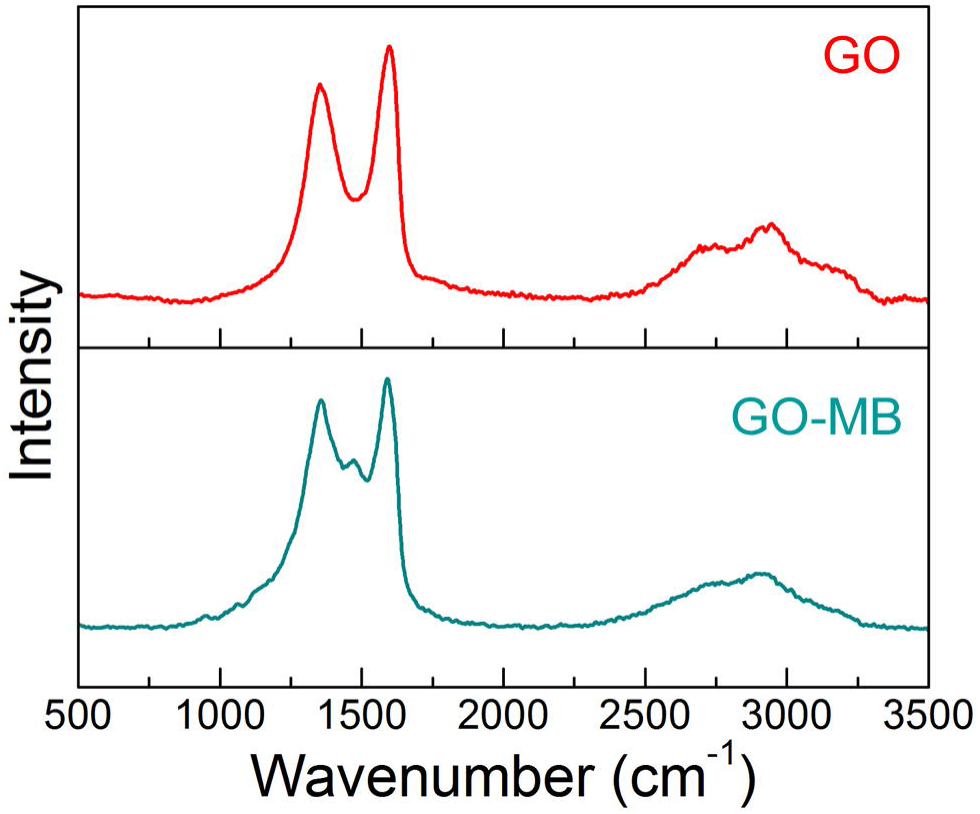
SEM image of (A) CSPE, GO- and GO□MB-modified electrodes.

Representative CVs measured during the process of electroadsorption of MB onto the surface of GO-modified electrode is provided in Fig. S4. A pair of electrochemically reversible redox peak is observed at □0.23 V (oxidation potential) and at □0.28 V (reduction potential). The constant increase of the redox peak currents with successive increase of scan cycles suggests that the electroadsorption of MB molecules on the surface of GO-modified electrode is indeed achieved. Observed redox behavior, due to MB ↔ leuco MB, is in close agreement with glassy carbon electrodes modified with other types of carbon nanostructures (viz., reduced GO, graphene and CNTs) adsorbed with MB [24,31,32]. In addition to electrostatic interaction, the mechanism behind the adsorption of MB on GO-based nanomaterials could also be possible through π-π stacking [24]. Further, increase of potential scans on GO-modified electrode might result in mild-to-moderate reduction of oxygen content, which could also mediate hydrophobic interactions with MB.

Fundamental structural changes occurred on the carbon lattice network of GO and MB-electroadsorbed GO nanosheets were studied from the Raman spectroscopy. Fig. 2 shows the Raman spectrum of GO modified SPE, which has two characteristic peaks at 1349 and 1597 cm^-1^ that attributed to the D- and G-bands [11], respectively. The intensity ratio of the D- and G-band peaks, *I*_(D)_/*I*_(G)_, of GO is 0.85. The overtone peaks of GO such as 2D band, D+G band and 2G band were observed at 2696 cm^-1^, 2925 cm^-1^ and 3160 cm^-1^, respectively [28]. After electroadsorption of MB, noticeble shift has been observed in the peak positions of D- and G-bands at 1355 and 1590 cm^-1^, respectively. Observed red shift in the G-band and increment in the *I*_(D)_/*I*_(G)_ ratio (0.92), indicates that the sp^2^ graphitic carbon region on the GO is significantly influenced by the electroadsorption of MB. The new moderate peak appeared at 1470 cm^-1^ is corresponded to the vibrational modes of MB, in agreement with thermodynamically prepared GO-MB nanocomposite [29]. Further, broadening of overtone peaks observed in the high wavenumber region of MB-electroadsorbed GO, suggested the structural changes in the carbon lattice and its associated layer properties. The average crystallite size of graphitic sp^2^ carbon domains in GO and MB-electroadsorbed GO modified electrodes, calculated according to modified Tuinstra-Koening equation [11,30] is 19.70 and 18.20 nm, respectively. These results suggest that the redox active MB molecules are successfully modified on the surface of GO nanosheets.

**Fig. 2.**
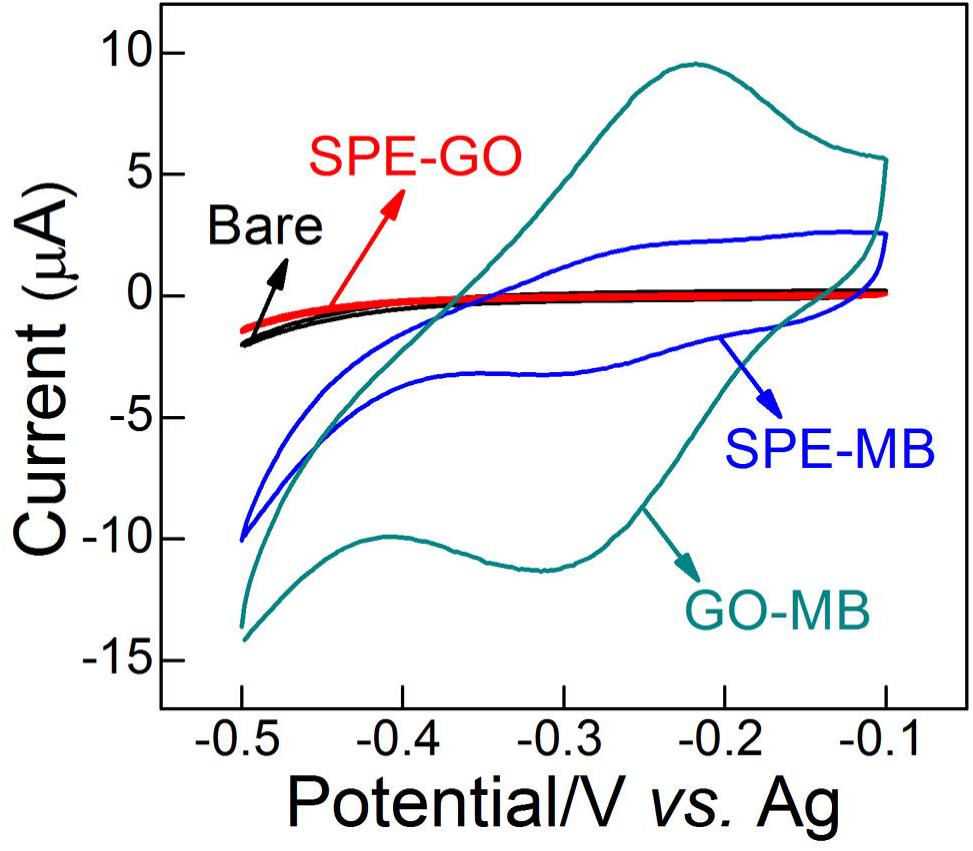
Raman spectral study on the surface of GO and MB-electroadorbed GO (GO-MB) nanosheets modified electrode.

### 3.3 Electrochemical properties of the modified electrodes

Fig. 3 displays the comparative CVs of SPE modified with different materials measured at a constant scan rate of 50 mV s^-1^, in presence of 10 mM PBS as supporting electrolyte. From the studied experimental conditions bare SPE and SPE □ GO are observed to have no distinct redox behavior. Although electroactive MB dye modified electrodes exhibit better redox behavior compared to bare SPE and SPE □ GO, because of its physical adsorption MB drop casted electrodes doesn’t enable durable redox behavior at the interface. On the other hand, the MB-electroadsorbed GO (i.e., GO□MB) electrode exhibit a reversible redox behavior with an anodic peak potential at □0.22 V and a cathodic peak potential at □0.31 V, which are attributed to the oxidation-reduction reaction of electroadsorbed MB [24,31,32]. The redox peak current intensities (*I*_pa_= 9.5 μA and *I*_pc_ = □11.27 μA) of this hybrid electrode material is observed to be superior than other studied electrode materials. In general, concentration of the redox molecules, nature of chemical groups exist on the electrode materials, scan rate, number of potential scan cycles and pH of the electrolyte solution are the crucial factors which determines the redox behavior of any redox molecules adsorbed onto the electrode surface. In this work, an optimal concentration of 50 μg mL^-1^ of MB was supplemented in the 10 mM PBS solution (pH 7.4), scan rate was 100 mV s^-1^ and the number potential scans were set to 20.

**Fig. 3.**
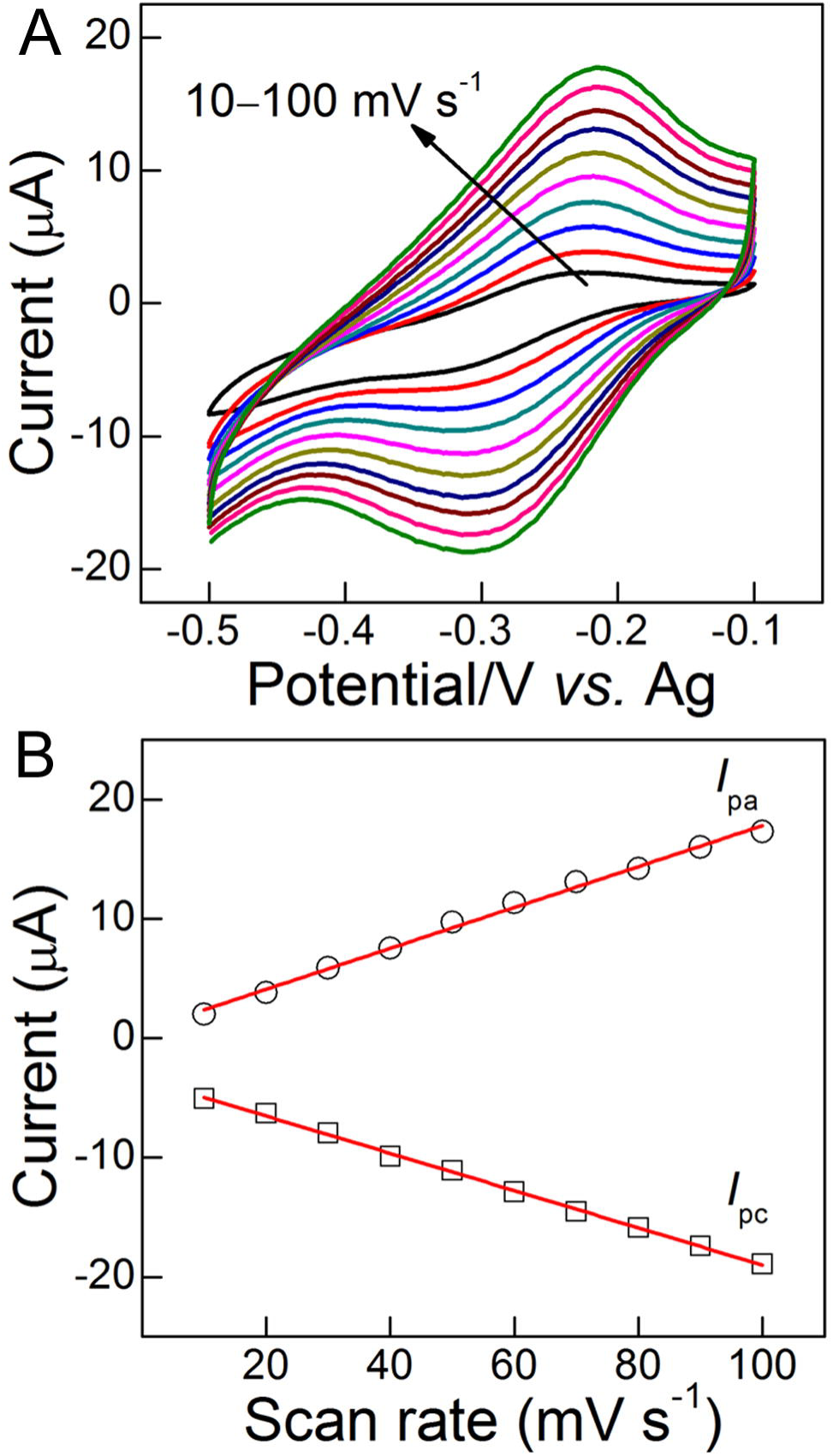
CVs of different materials modified electrodes recorded in 10 mM PBS (pH 7.4), at a scan rate of 50 mV s^-1^.

From the complementary studies discussed in the previous section (surface morphology, elemental analysis and Raman spectroscopy), it is indeed clear that the continuous larger carbon lattice network of GO nanosheets with abundant hydrophilic oxygenated functional groups are exceptionally useful in the electroadsorption of MB. It is expected that the observed durable and amplified redox behavior of GO□MB modified electrodes could be suitable for biosensor application. Fig. 4A depicts the CVs of MB-electroadsorbed GO modified electrode in PBS at various scan rates in the range 10 to 100 mV s^-1^. As shown, the anodic peak currents linearly increase with the scan rates while the cathodic peak current linearly decrease with the scan rates. Observed pair of well-defined and broad redox wave without appreciable change in the peak potential demonstrates a fast electron transfer process. Corresponding linear fit for the anodic and cathodic peak currents with the scan rates provided a correlation co-efficient of 0.9950 (*I*_pa_) and 0.9988 (*I*_pc_), respectively. These suggest that the generated redox behavior from the MB-electroadsorbed GO nanostructure on SPE is a surface-confined process, which are promising for the biosensor platform design.

**Fig. 4.**
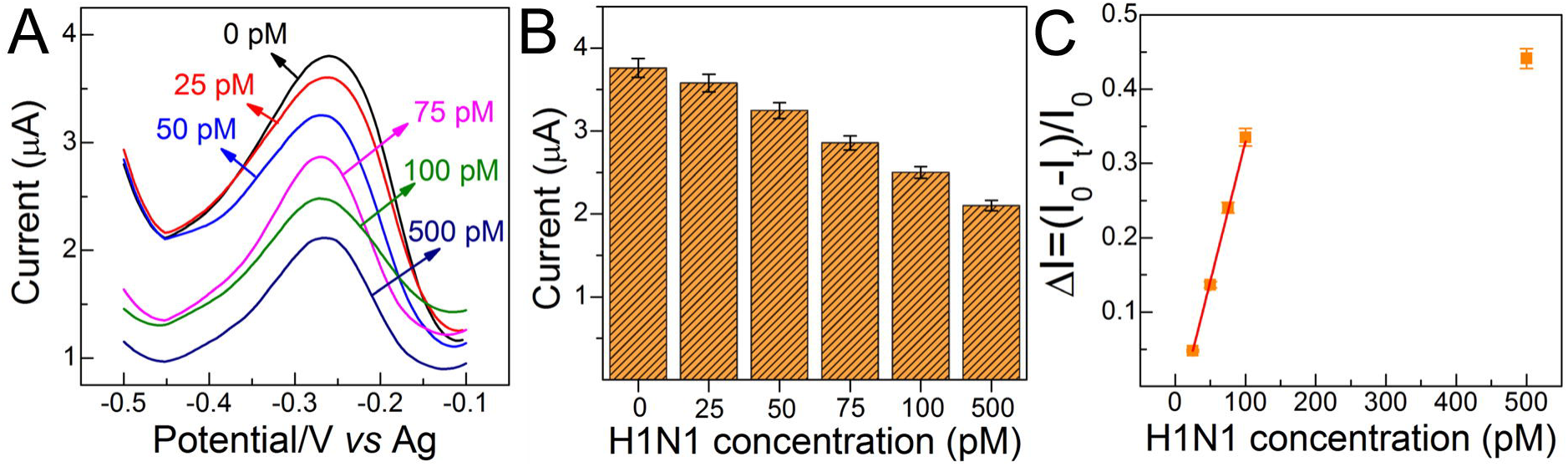
(A) CVs of GO-MB modified electrodes at different scan rates (10-100 mV s^-1^) in 10 mM PBS (pH 7.4). (B) The plots of anodic and cathodic peak currents versus scan rate.

### 3.4 Immunosensing of H5N1 and H1N1

As described in the experimental section 2.4, bio-active layers based on CS and protein-A molecules were modified on the surface of sensor element (GO□MB) to enrich the immobilization and performance of mAb. As shown in Scheme 1, two working substrates (WE1 and WE2) were suitably modified with different mAb such as mouse anti influenza-A hemagglutinin H5N1 mAb and mouse anti influenza-A hemagglutinin H1N1 mAb. Unlike conventional single working electrodes, dual SPE chip is advantageous in terms of simultaneous detection of two different analytes in the same sample, at the same time. Measurement of electrochemical signals and data processing can be performed sequentially. Upon the specific interaction of target antigens onto the surface of mAb-modified sensor element an insulating layer could be formed, due to the immune complex. Such binding creates a barrier toward the inherent electron-transfer process at the electrode interface. This altered electrochemical process can be correlated to the concentrations of the target analyte. For the proof-of-concept, two recombinant protein samples were utilized in this study, viz., hemagglutinin influenza-A virus H5N1 Vietnam 1203/04 and hemagglutinin influenza-A virus H1N1 California 04/2009. Based on the well-known redox behavior of the GO□MB electrodes in PBS buffer the DPV study was performed in the potential region □0.5 to □0.1 V, at a scan rate of 50 mV s^-1^ with an amplitude of 2 mV and a step potential of 4 mV.

Fig. 5A depicts the representative DPVs of immunosensor GO□MB/CS/protein-A/anti-H1N1 electrodes studied with different concentrations (0, 25, 50, 75, 100 and 500 pM) of antigen H1N1 diluted in PBS. As can be seen, the inherent anodic peak (*E*_pa_ = □0.26 V, attributed to redox reaction of MB-electroadsorbed GO) current response from the immunoelectrode interacted with 500 pM concentration of H1N1 exhibited the least peak current intensity (2.1 ± 0.06 μA pM^-1^) than other studied samples, suggesting an high ratio of immune complex formation on the sensor surface. With different analyte concentration a distinguishable changes in the anodic peak current were clearly observed, as revealed in the histogram plot (Fig. 5B). A linear fit according to the DPV results expressed in Δ*I* is shown in Fig. 5C, Δ*I* = (*I*_0_□*I*_*t*_/*I*_0_), where *I*_0_ is the absolute peak current value of the modified electrode tested in PBS without antigens and *I*_t_ is the absolute peak current value of the modified electrode measured in PBS with the desired concentration of an antigen. The regression equation for the experimental data was *y* = *A* + *B*(*X*), where *y* is the sensor response measured as the change in current Δ*I*, *X* is the concentration of analyte, and *A* and *B* are the sensor constants. Linear fitting for the data points in the Fig. 5C gave *y* = □0.04622 + 0.00376 (*X*) with a correlation co-efficient of 0.9978. From the studied, DPV technique and, analyte (H1N1) concentration range the detection limit has been determined according to the 3*γ*/*m* criteria, where *σ* represents the standard deviation of blank and *m* denotes the slope of the calibration plot and was found to be 9.4 pM.

**Fig. 5.**
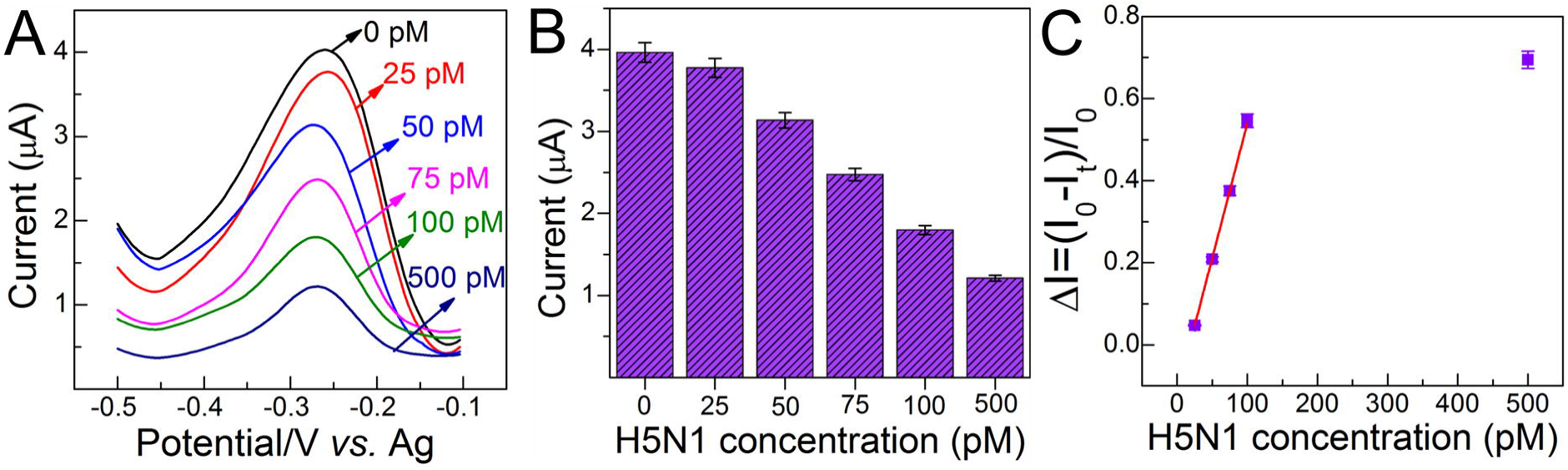
(A) Representative DPVs of GO-MB/CS/protein-A/anti-H1N1 electrodes against various concentrations of antigen H1N1 and (B) the selected relevant peak voltammetric histogram. (C) Calibration curve derived from the DPVs expressed in ΔI. ΔI=(I_0_□I_t_/I_0_), where I_0_ is the peak current value of modified electrode in PBS without antigen application and I_t_ is the peak current value of the modified electrode measured in PBS with the desired concentration of an antigen (*n*=3).

Similarly, the DPVs recorded from WE2, the immunosensor GO□MB/CS/protein-A/anti-H5N1 tested with different concentrations of H5N1 antigens in PBS is shown in Fig. 6A. Due to its specific affinity mAb H5N1 modified sensor platform exhibited a well-resolved correlation between the change of current and analyte concentration. As represented in the histogram plot (Fig. 6B) among the studied concentrations, the immunoelectrode treated with 500 pM sample showed the lowest peak current intesnity 1.21 ± 0.03 μA pM^-1^, indicating a concentration depended linear voltammetric response. Linear fitting for the data points in the Fig. 6C gave a regression equation *y* = □0.11715 + 0.00657 (*X*) with an R^2^ = 0.9997. The detection limit studied (according to 3*σ*/*m*) in the present experimental concentration range was found to be 8.3 pM.

**Fig. 6.**
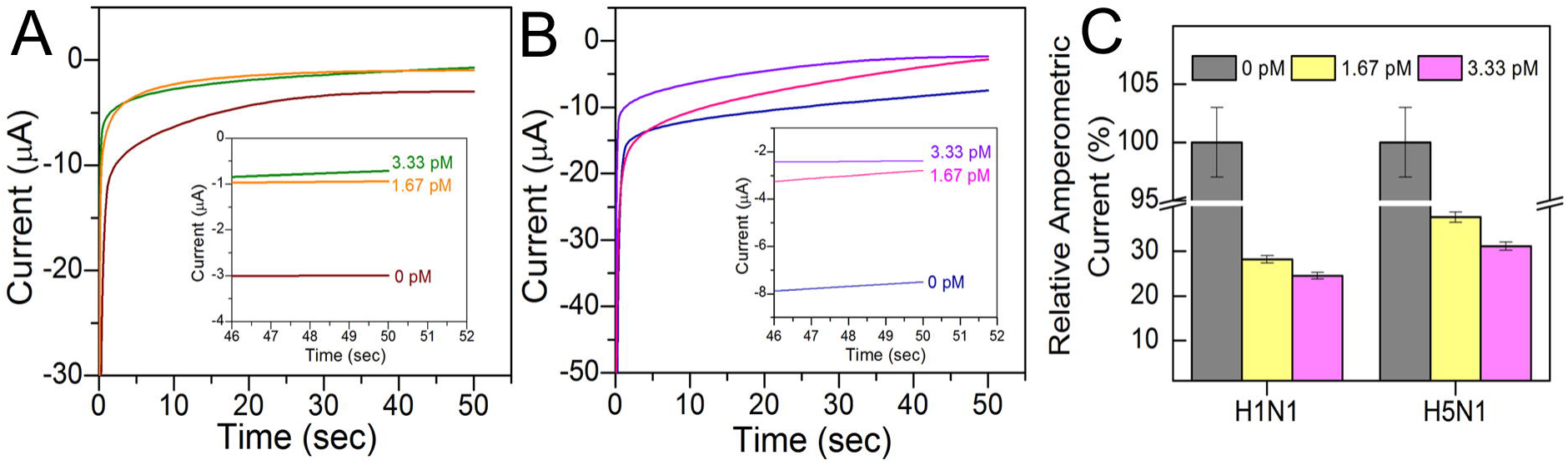
(A) Representative DPVs of GO-MB/CS/protein-A/anti-H5N1 electrodes against various concentrations of antigen H5N1 and (B) the selected relevant peak voltammetric histogram. (C) Calibration curve derived from the DPVs expressed in ΔI (*n*=3).

In order to explore a further feasibility of the proposed immunosensing concept in this work a different customized single electrode chip were utilized. Single carbon SPE integrated with carbon counter and Ag/AgCl reference electrode, were procured from Pine research instrumentation, NC, USA. Similar to the dual electrodes, fabrication of sensor element and sequential modification of bioactive layers and antibodies were achieved. Based on the preliminary voltammetric study it was found that an optimal potential of □0.275 V (responsible for redox reaction of GO□MB) is appropriate for the chronoamperometric study. Fig. 7(A) and (B) shows the representative chronoamperometric current signals based on the immobilized antibodies (mAb H1N1 and mAb H5N1) and respective antigens of different concentration. As can be seen both the immunosensor platform interacted with higher concentration of antigen samples (3.33 pM) exhibited the lowest reduction peak current, □7.11 μA (H1N1) and □2.38 μA (H5N1), in complement with DPVs. Provided inset figure clearly reveals the differential current signals with the studied antigen concentrations (0, 1.67 and 3.3 pM). As the redox behavior of MB-electroadsorbed GO electrode (Fig. 3, GO□MB trace) exhibited a near unity of the ratio of *I*_pa_/I_pc_, from the applied voltage □0.275 V (i.e., cathodic peak potential, *E*_pc_) the immunoelectrodes showed the absolute reduction current signals. For better clarity, a relative chronoamperometric signals are determined and expressed in the histogram (Fig. 7C). Obtained chronoamperometric results certainly support that the developed immunosensor platform enabled a satisfactory specificity toward the different AI viral strains with a picomolar sensitivity. The prototype detection approach demonstrated from the chronoamperometry is comparatively faster in analytical response time, <1 min than most of the conventional molecular methods. Furthermore, unlike recent other electrochemical immunosensor designs [12,13,33,34], the proposed immunosensor platform didn’t require additional external redox marker like [Fe(CN)_6_]^3-^/^4-^ to monitor the signals, this is because of the inherent redox-activity of GO□MB sensor element. Demonstrated dual sensor design would not only provide rapid analysis and cost-efficiency in assay but also create feasibility toward simultaneous detection of different pathogens in the same sample.

**Fig. 7.**
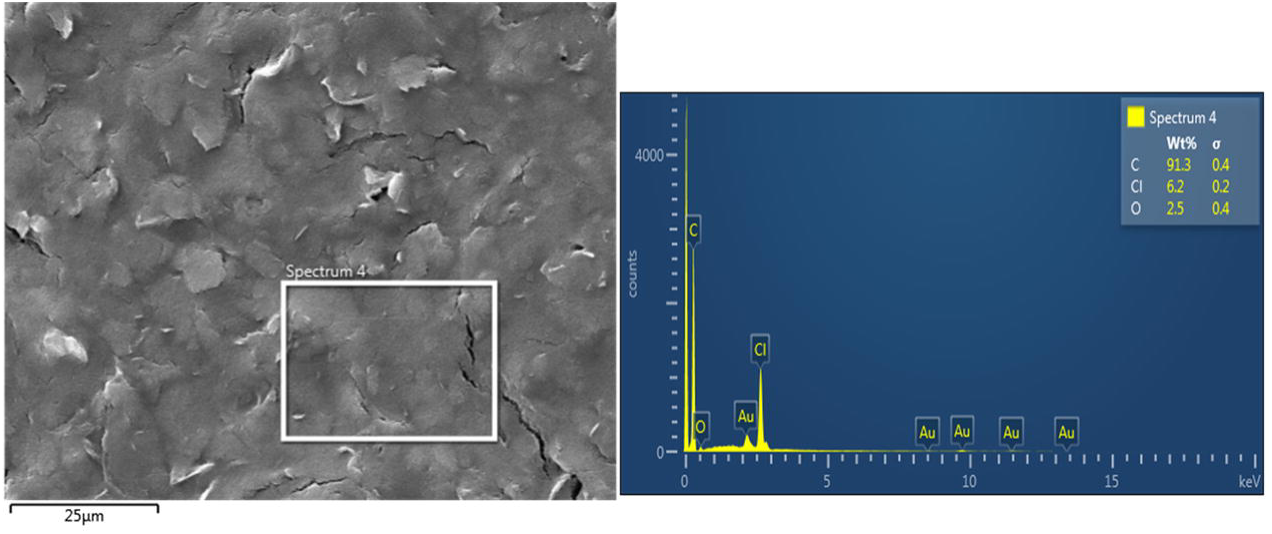
(A) Chronoamperometric curve recorded from GO-MB/CS-protein-A/anti H1N1 *vs* different concentrations of antigen H1N1. (B) Chronoamperometric curve recorded from GO-MB/CS-protein-A/anti H5N1 *vs* different concentrations of antigen H5N1. Inset shows the selected zone of chronoamperometric curve. (C) Relative amperometric histogram represents % difference observed from the modified electrodes at three different concentrations (*n*=3).

## 4. Conclusions

A dual sensor platform was designed, based on electrochemical property of MB-electroadsorbed GO nanostructure and immunoaffinity of immobilized mAb. Functionalization of bio-active molecules (chitosan and protein-A) at the interface of sensor element significantly amplified the immobilization of mAb, which eventually supported the durable immune complex formation with target antigens. The voltammetric and chronoamperometric signals generated from the proposed immunoelectrodes are independent due to its in-built redox-reaction at the interface, without the need for external electro-active marker. In the presence of selective mAb the two sensor substrates enabled the simultaneous detection of different subtypes of influenza-A virus antigen. The designed dual immunosensor is facile in construction and offers high sensitivity (picomolar level), rapid analysis time (<1 min) and reproducibility. Considering the demands for the rapid detection of influenza threats in poultry facilities and associated fatal complications, a simple and sensitive influenza assay is quite beneficial. Further optimization of the developed immunosensor design could be escalated for the multiplexed detection and discrimination of various high and low-pathogenic influenza viral strains.

## Acknowledgments

The authors sincerely thank the Natural Sciences and Engineering Research Council of Canada, Canadian Poultry Research Council, Mitacs Accelerate Fellowship program and Egg Farmers of Canada for funding this study.

## Appendix A. Supporting information

Supplementary data associated with this article can be found in the online version.

## Figure Captions

**Fig. S1.**
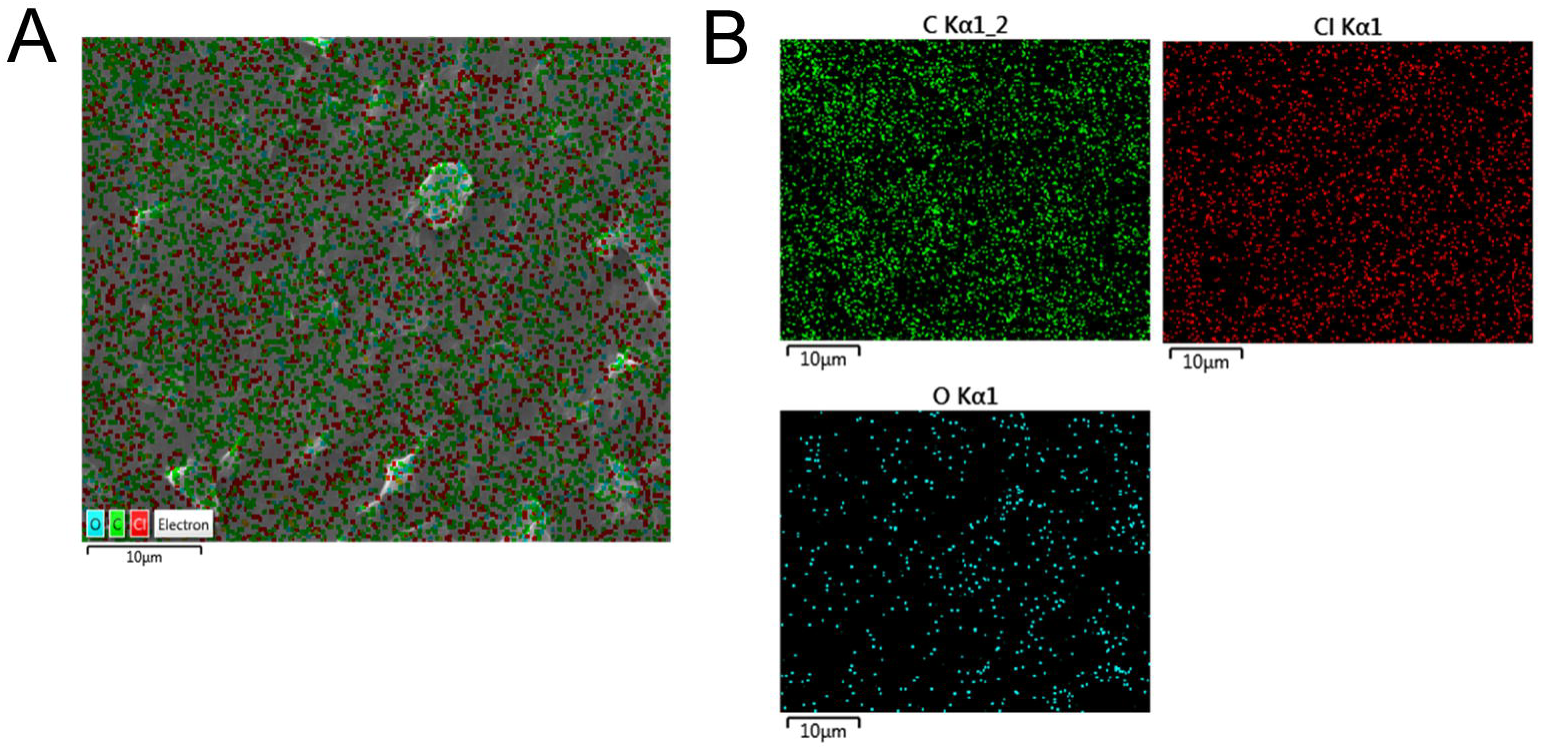

**Fig. S2.**
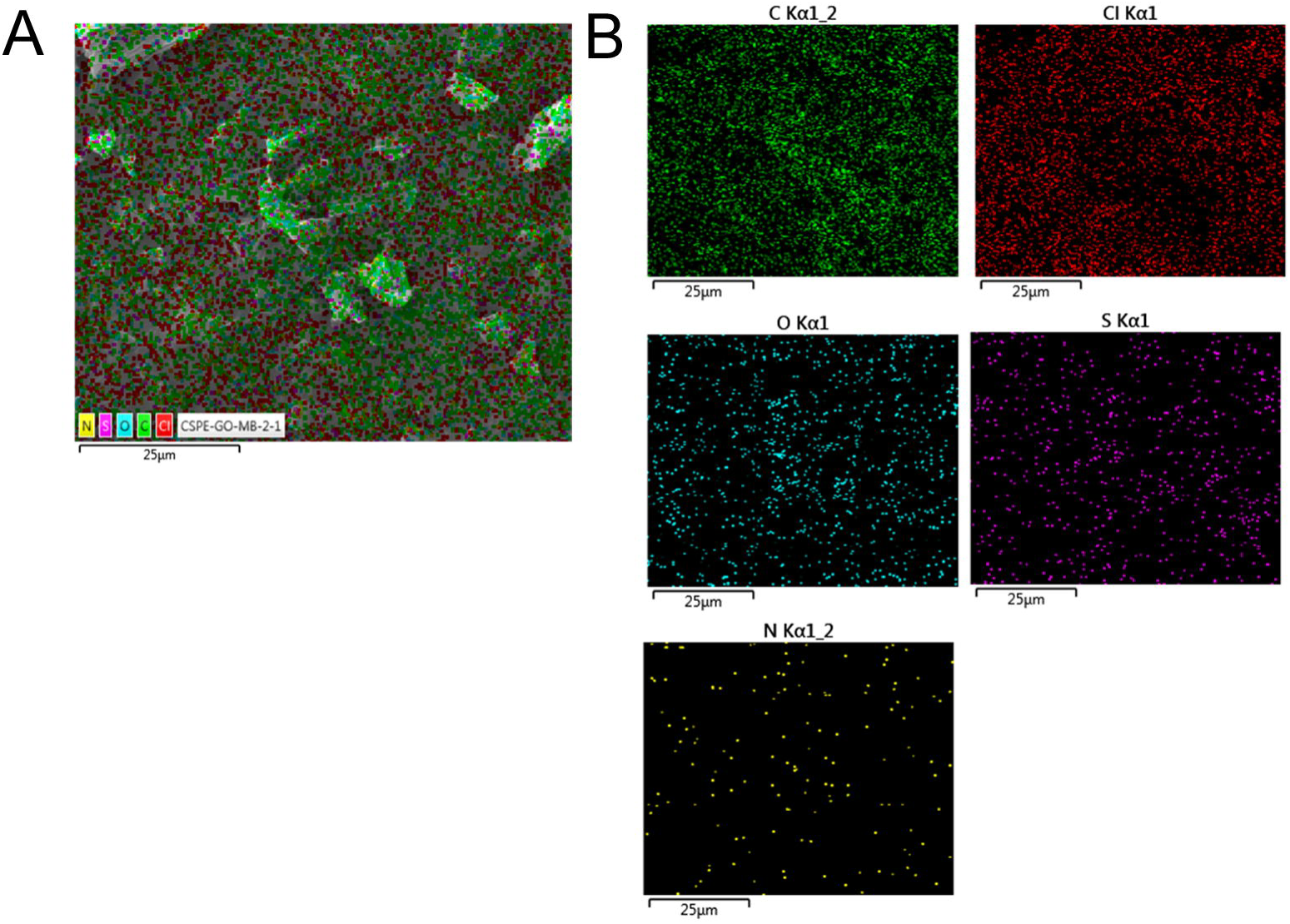

**Fig. S3.**
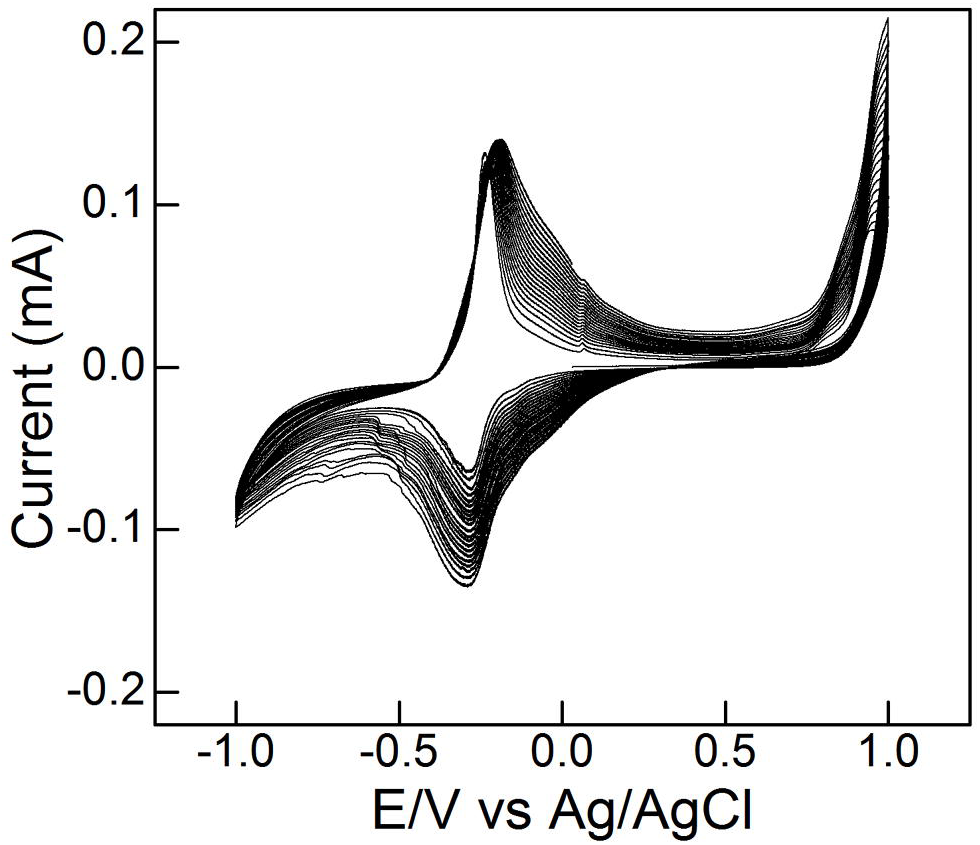

**Fig. S4.**
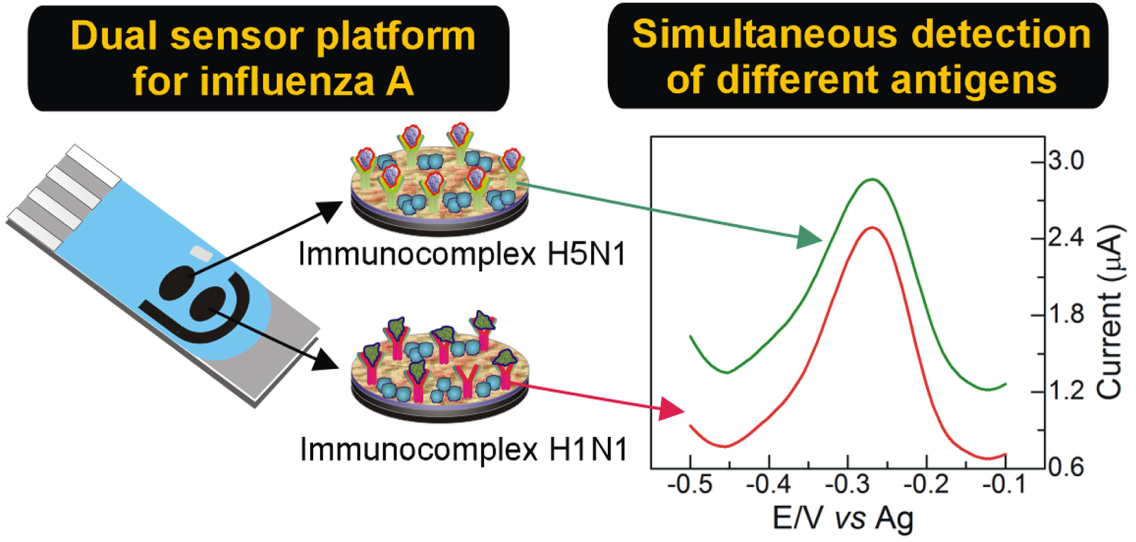

**Fig. S5.**
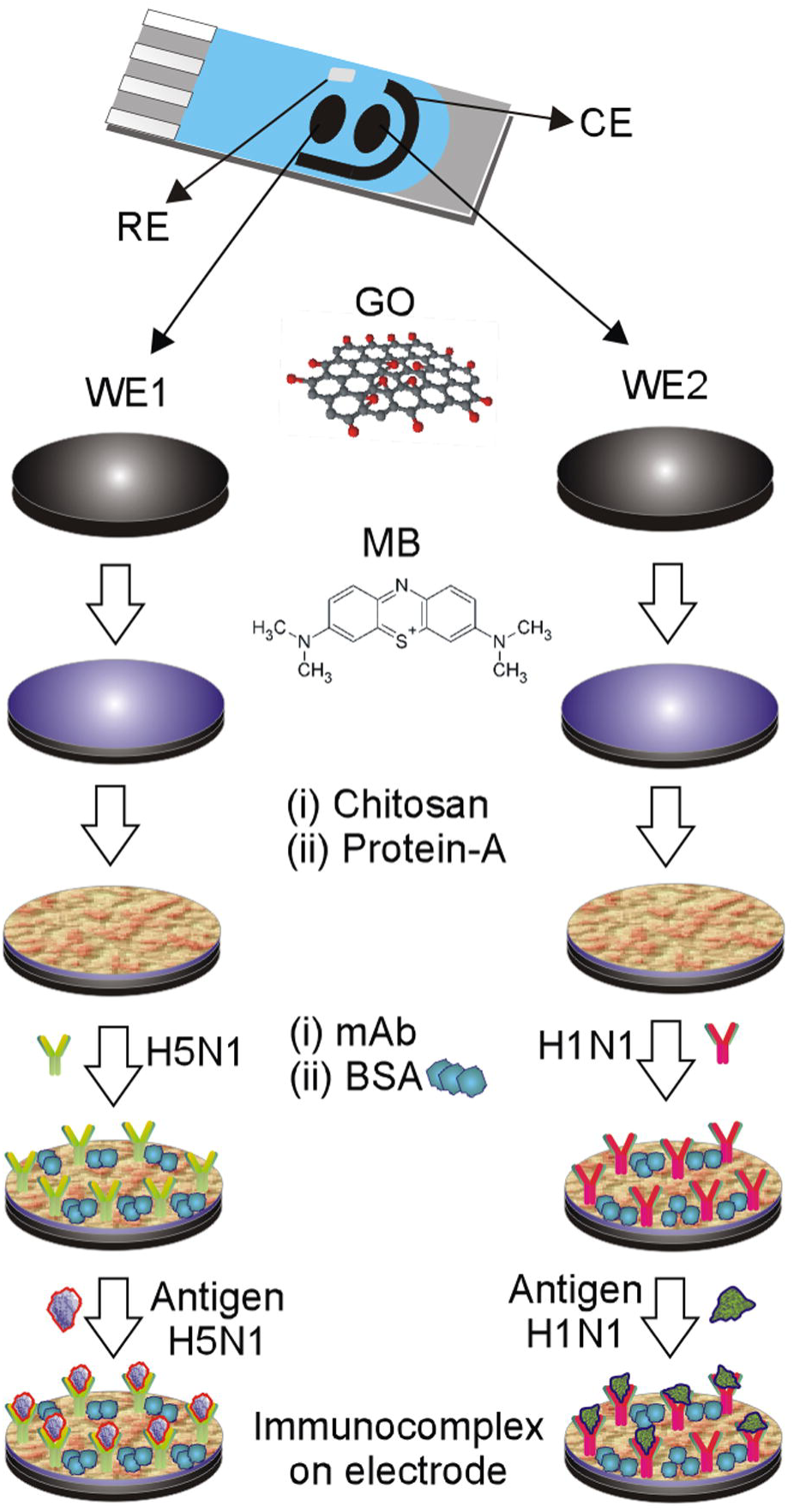

